# Gradients of orientation, composition and hydration of proteins for efficient light collection by the cornea of the horseshoe crab

**DOI:** 10.1101/2022.02.21.481013

**Authors:** Oliver Spaeker, Gavin Taylor, Bodo Wilts, Tomáš Slabý, Mohamed Ashraf Khalil Abdel-Rahman, Ernesto Scoppola, Clemens NZ Schmitt, Michael Sztucki, Jiliang Liu, Luca Bertinetti, Wolfgang Wagermaier, Gerhard Scholtz, Peter Fratzl, Yael Politi

## Abstract

The lateral eyes of the horseshoe crab, *Limulus polyphemus*, are the largest compound eyes within recent Arthropoda. While this visual system has been extensively described before, the precise mechanism allowing vision has remained controversial. Correlating quantitative refractive index (RI) mapping and detailed structural analysis, we demonstrate how gradients of RI in the cornea result from the hierarchical organization of chitin-protein fibers, heterogeneity in protein composition and bromine doping, as well as spatial variation in water content. Combining the realistic cornea structure and measured RI gradients with full-wave optical modelling and ray-tracing approaches, we show that the light collection mechanism depends on both refraction-based graded index (GRIN) optics and total internal reflection. The optical properties of the cornea are governed by different mechanisms at different hierarchical levels, demonstrating the remarkable versatility of arthropod cuticle.

**One-sentence summary:** Structural hierarchy and protein hydration determine the optical performance of the cornea of *L. polyphemus*.

## Body

Arthropod vision has fascinated researchers for more than a century for its diverse optical mechanisms (*1*) and recently for potential bio-inspired design of micro-lens-arrays (*2*). Their compound lateral eyes form an array of individual light collecting and sensing units, the ommatidia, typically consisting of a cornea, a conical lens or light guiding device, and a receptor unit called rhabdom (*3*). Light entering the ommatidium is guided towards the rhabdom by refraction, reflection or a combination thereof (*4*). An image is created by the integration of stimuli from each receptor unit, which receives the light from optically isolated ommatidia in apposition eye systems, or multiple neighboring ommatidia in superposition systems (*3*).

Vision is especially well studied in the Atlantic horseshoe crab, *Limulus polyphemus* (*L*. 1758, *Xiphosura*) (*5*). Horseshoe crabs have six rudimentary eyes, two ocelli and a pair of lateral compound eyes (Fig. 1A) (*6*). While the latter are very well understood from a neurophysiological perspective (*7*), a debate remains as to their light focusing mechanisms (*8*). Exner (*1*) proposed and Land (*8*) measured a radial nearly parabolic refractive index (RI) gradient in the cones leading to a refraction-based focusing mechanisms, termed cylinder-lens by Exner (*1*) and described today as graded Index (GRIN) optics. Conversely, Levi-Setti et al. (*9*) emphasized the cone shape and ascribed the focusing power of the lens to total internal reflection (TIR) at the interface between cornea and surrounding tissue, suggesting it was an example of an optimal light collector. Although the shape of the cones and the general radial RI gradient in the lenses are well described, to date little is known about the structure-property-function relationships in the material that forms the lens. Here we determine how the optical properties of the lens stem from its multi-scale architecture and local composition. To that end, we have mapped the RI along and across the cones and correlated the results with spatially resolved structural and compositional analyses of the material. Using different optical modeling approaches, we demonstrate that both the observed GRIN profile and lens shape have roles in determining the overall optical behavior.

**Figure 1:**
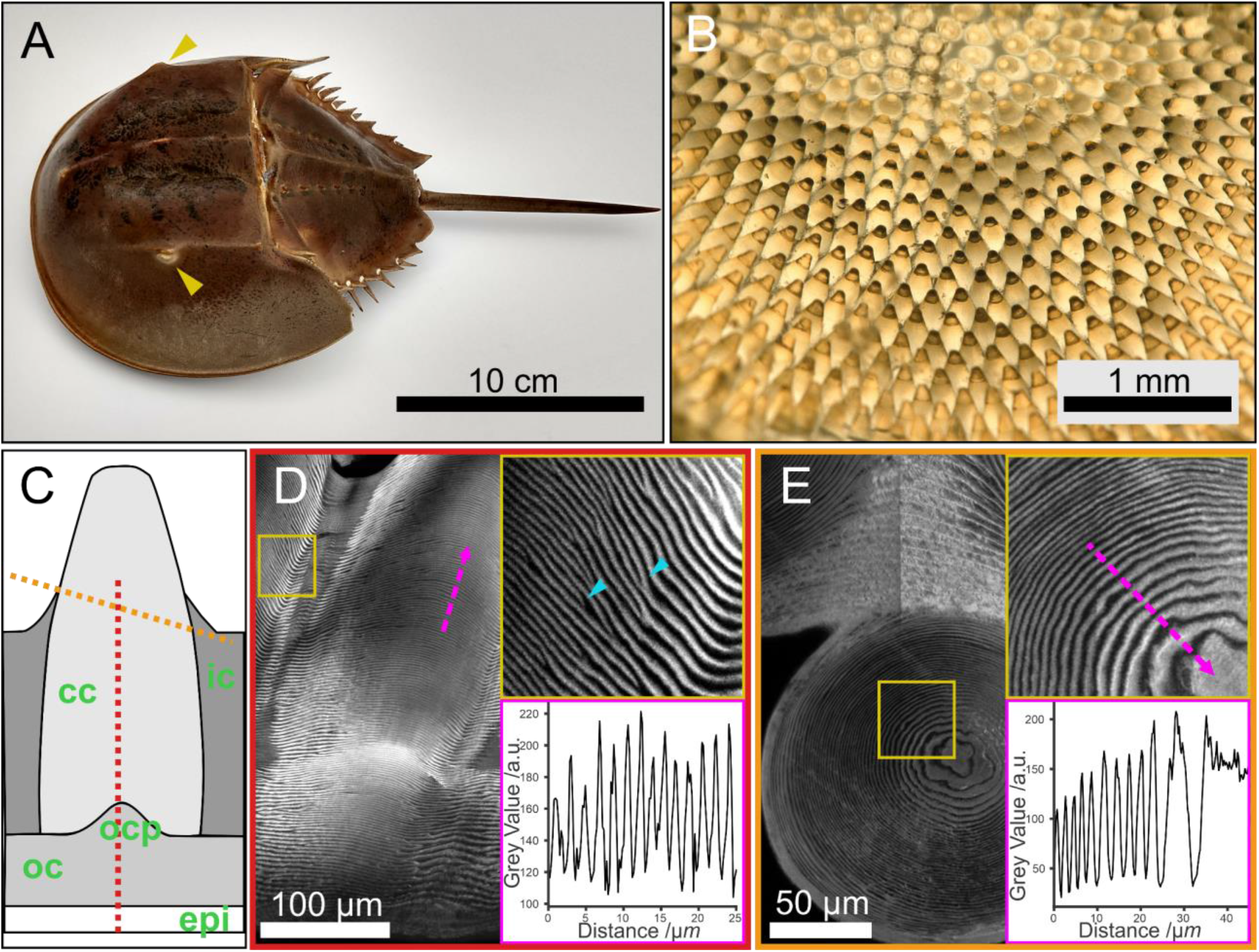
Structure of *L. polyphemus* cornea. **(A)** Image of *L. polyphemus* showing the position of the lateral compound eyes (yellow arrowheads) **(B)** Transmission light micrograph taken from the proximal side of a cornea after removal of cells and pigments. Note the dark and bright appearance of the cone depending on the viewing angle. **(C)** Schematic depiction of corneal cones (cc) inllustrating the different regions discussed in the text, intercone (ic), epicuticle (epi), outer cornea (oc), outer-cornea protrusion (ocp). The yellow and red dotted lines indicate the section orientation in D and E. **(D)** A longitudinal section (red dotted line in C) stained with DY96 and imaged with CLSM readily reveals the helicoidal arrangement of chitin in the *L. polyphemus* cornea, as can be deduced from the sinusoidal alternation of light and dark bands. Upper inset: magnification of the cone-intercone border, yellow square, showing defects in lamella organization (blue arrowheads). Bottom inset: variation in grey level along the magenta dotted arrow. A section of a full cone shown in Fig. S1B **(E)** CLSM of DY96-stained cross sections (orange dotted line in C). Upper inset: magnification of the cone-centre, marked in yellow square. Bottom inset: variation in grey level along the magenta dotted arrow. The bright and dark bands broaden towards the centre due to lamellae inclination. The brightness difference around the midline is a stitching artefact.

The *L. polyphemus* lateral eye consists of an array of lenses forming a cornea and inwards projecting cones (Fig. 1B and C, Fig. S1). In contrast to the situation in most insects and crustaceans and presumably representing the ancestral condition for arthropod eyes (*10*), the cones are formed by the cuticle, the same material that builds the animal’s exoskeleton. Arthropod cuticle is made of a chitin-protein fiber composite that shows structural and compositional versatility which allows its multifunctional roles (*11*). In *L. polyphemus*, the corneal cones are hierarchically structured: the chitin-protein fibers are organized helicoidally leading to a lamellated appearance, where the lamellae (i.e. fiber-sheet layer consisting of half helicoidal pitch) organize in a curved nested arrangement that follows the cone geometry (Fig. 1D and E). Image stacks acquired by confocal laser scanning microscopy (CLSM) of Direct Yellow 96 (DY96)- stained sections of the cornea reveal a regular 3.52 μm ± 0.49 μm helicoidal pitch across and along the cone, with radially increasing lamella-inclination from the center to the edge of the cone (Fig. 1E). Here, the cones are embedded in a cuticular region, hereafter termed intercone, showing an increased helicoidal pitch of 5.96 μm ± 1.04 μm. The mismatch in helicoidal pitch dimensions leads to multiple defects, especially concentrated at the interface region between cones and intercone (Fig. 1D arrowheads). At the distal side, the cones are connected to the outer-cornea through a structure that we termed the outer-cornea protrusion, and a thick epicuticle constitutes the interface with the environment (Fig. 1C and D). The arthropod cuticle is typically perforated by multiple pore-canals that serve for transport across the cuticle. In *L. polyphemus* compound eyes, pore-canals are only present in the intercone but not within the cones or other regions of the cornea (Fig. S1C).

We investigated the hierarchical structure of the corneal cones at the μm- and nm-scale by scanning X-ray diffraction (XRD) using cornea thin sections in the hydrated and dry state (Fig. 2). The acquired XRD profiles show the typical reflections of α-chitin with the (020) reflection at *q* ~ 6.4 nm^-1^ to 6.6 nm^-1^, (110) at *q* = 13.6 nm^-1^ and the (013) at *q* = 18.5 nm^-1^ (Fig. 2A) (*12–14*). Additionally, in the small-angle scattering (SAXS) region, a correlation peak resulting from chitin-protein fiber packing (*15*), is observed at *q* ~ 1.1 nm^-1^. Scanning the sample using an X-ray beam with 300 nm cross-section reveals heterogeneity in the cornea’s material composition and structure. Figure 2Ai shows XRD profiles of various regions within the cornea. However, at the outer-cornea surface, chitin reflections are absent and instead the diffraction profile shows two broad peaks at *q* = 6 nm^-1^ and q = 14 nm^-1^ (Fig. 2A, violet curve). The occurrence of these peaks is indicative of cross-beta structures in proteins (*16*, *17*). Cross-beta protein organization is common in arthropod cuticles (*18*, *19*), however it is often difficult to discriminate the protein reflections from those of chitin as they typically coincide and since chitin-poor or protein-rich layers are usually thin with respect to the probed volume. In *L. polyphemus* an especially thick epicuticle (the part of the cuticle that is devoid of chitin) allows direct insight into the dominant protein structure in the cuticle.

**Figure 2:**
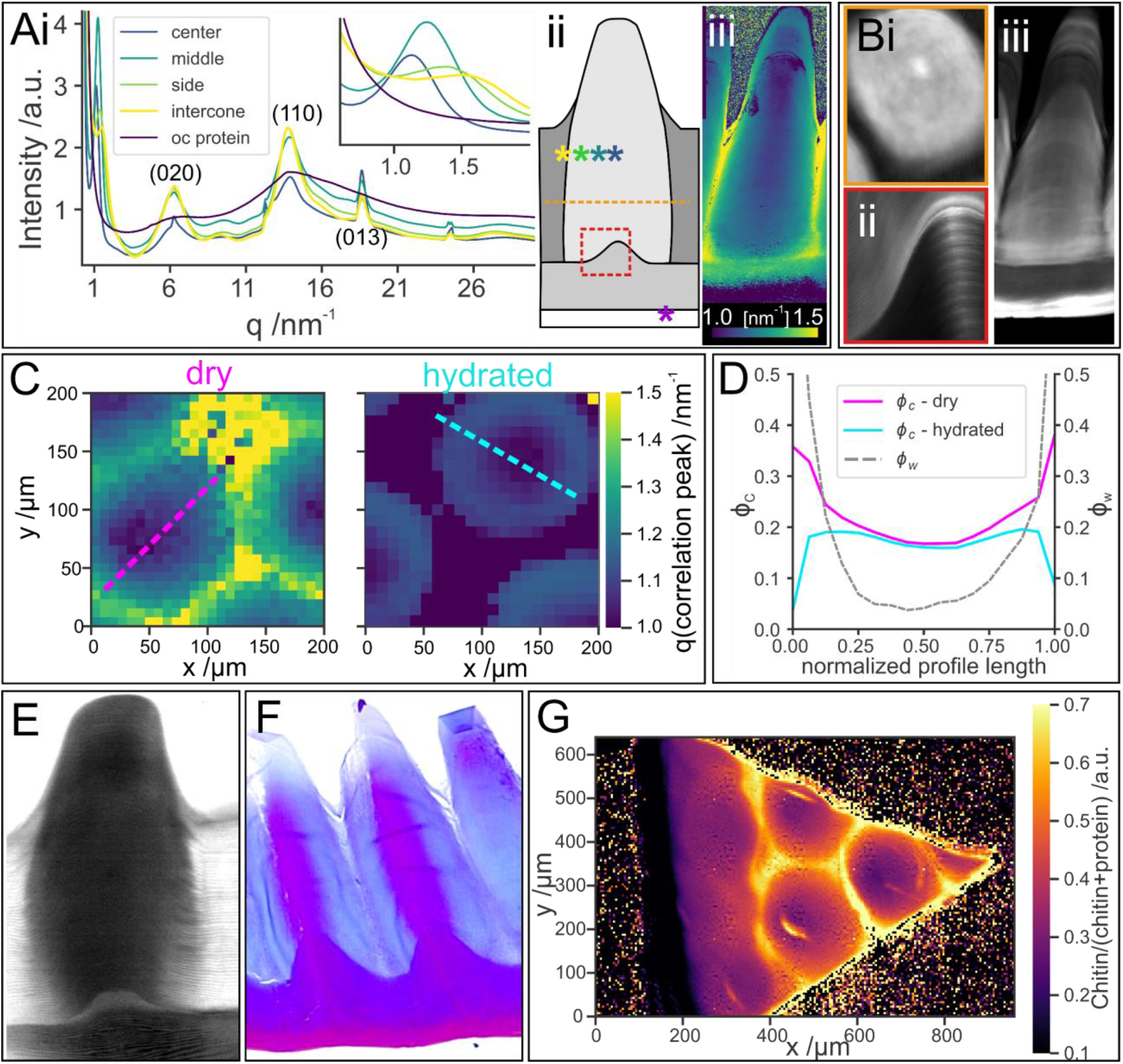
Structure and compositional variations along the cornea. **(A)** XRD mapping (i) Representative XRD profiles obtained by averaging diffraction patterns at different regions of the *L. polyphemus* cornea measured dry. The inset shows a magnification of the chitin correlation peak around 1.1 nm^-1^. The positions within the cornea are indicated by color-coded asterisks in the cornea scheme in A(ii), which also contains positions for B(i) and B(ii). A(iii) Mapping of the correlation peak *q*-position across a dried longitudinal section. **(B)** Br-XRF maps of the cornea (i) High resolution XRF maps of BrKα of a cone cross section and (ii) a longitudinal section at the outer cornea protrusion and (iii) a longitudinal section of the entire cone. The section is cut off axis therefore it does not contain the outer cornea protrusion. **(C)** The effect of hydration on the *q*-position of the fiber correlation peak in cornea cross sections. The central values are similar in both samples, whereas towards the edges of the cones they shift to lower *q* in hydrated (right) relative to dry (left) samples, indicating that swelling due to hydration is limited to this region. Dotted lines indicate the positions of the profiles shown in D **(D)** Chitin and water volume fractions calculated from the correlation peak position extracted from maps in C (see eq. 1). The dimensions of the cones are normalized to accommodate for the different dimensions between dry and hydrated state. **(E)** Raman hydration map (integrated intensity of the OH bands between 3111 cm^-1^ and 3689 cm^-1^) **(F)** Mallory trichrome staining of a longitudinal cornea section **(G)** FTIR map of a dry oblique section of the cornea containing the outer cornea. The map depicts the ratio of the chitin and protein signals as determined from the intensity ratio in the regions: 1700 cm^-1^ to 1600 cm^-1^ and 1180 cm^-1^ to 1000 cm^-1^, Fig. S3C.

In the SAXS region, the fiber correlation peak position shows large spatial variation (Fig. 2Ai, inset). In air-dried sections, the radial *q-*position of this peak changes from 1.1 nm^-1^ to 1.42 nm^-1^ within the cones and to 1.5 nm^-1^ in the intercone region (Fig. 2Aiii), corresponding to a shift in fiber-to-fiber distance from around 5.7 nm to 4.6 nm and 4.2 nm in real space, respectively. We estimated the chitin volume fraction (*φ_c,dry_*) across the sample assuming constant chitin crystallites diameter (see note in SI) and a hexagonal packing, using (*20*):

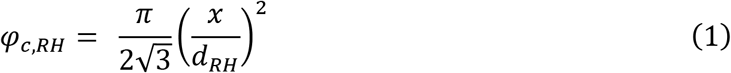

where *x* is the chitin crystal diameter (2.84 nm) and *d_RH_* is the fiber-to-fiber distance for either dry or hydrated samples retrieved from the position of the SAXS peak. The chitin volume fraction in dry samples changes from approximately 20 % in the cone center to 37 % at the cone border (Fig. 2D), where protein volume fraction, *φ_p,dry_*, is assumed as 1- *φ_c,dry_*. The variation in the chitin to protein ratio is also reflected in FTIR, which also demonstrates the absence of chitin in the outer layers of the cornea (Fig. 2G, Fig. S3C). In its natural environment the *L. polyphemus* cornea is fully hydrated. Hydrated cornea sections show an interesting swelling behavior; while the correlation peak shifts to lower *q* values at the sides of the cone in hydrated compared to dry cornea sections, indicating an increased distance between chitin fibrils, the *q* values measured in the center remain unchanged. The water volume fraction *φ_w_* was calculated using *φ_c,dry_, φ_p,dry_* and the chitin volume fraction in hydrated cornea cross sections, according to the equation:

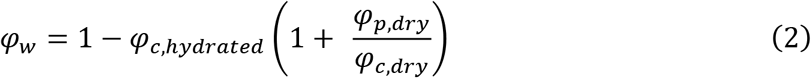

The water volume fraction shows a roughly parabolic profile with its minimum close to the cone center (Fig. 2D). This hydration gradient can also be visualized using micro-Raman spectroscopy by mapping the intensity of the OH bands between 3111 cm^-1^ to 3689 cm^-1^ (*21*) (Fig. 2E and S3D).

Cuticle hydration is tightly correlated with its sclerotization level, i.e. the incorporation of catechol derivatives that crosslink the matrix and render it hydrophobic, although the molecular details of these effects are not completely clear (*22*). To visualize the sclerotization level within the cornea, we used Mallory staining. This histological triple stain is sensitive to sclerotization and is typically used to differentiate between cuticle layers: unsclerotized cuticle (e.g. endocuticle) is stained blue, whereas sclerotization, as found in exocuticle and muscle attachment sites, leads to intense magenta coloration (*23*). Cones stained with Mallory display strong magenta coloration in their core and a blue-stained circumference indicating a cross-linked core and poorly sclerotized shell (Fig. 2F), in striking agreement with the hydration results from Raman and XRD.

Concurrently with XRD, we also recorded the emitted X-ray fluorescence (XRF) (Fig. 2B). We detected an elevated Br signal in the epicuticle, as previously observed in insects (*18*), but not in the outer cornea. Br was also present in the cones but not in the intercone region (Fig. 2Biii, Fig. S2Bii). In addition, a thin cuticular layer enriched with Zn surrounds the surface of the cones in the region where they extend out of the intercone layer (Fig. S2Ci). Interestingly, the Br distribution does not closely follow the sclerotization pattern determined by Mallory staining, suggesting that in *L. polyphemus* halogen incorporation is not coupled to the sclerotization process as suggested for insects (*24*).

To determine the contributions of the observed structural and compositional heterogeneities to the optical properties of the eyes, we performed spatial mapping of the RI (at 650 nm) in longitudinal and cross sections of the cornea using quantitative phase imaging (QPI) (*25*). This method allows quantitative determination of a phase shift of light transmitted through the sample. Using QPI acquired in the presence of two media with different RI (phase decoupling, *n_m1_* = 1.425, *n_m2_* = 1.457) (*26*), a spatially-resolved RI map can be calculated with μm-resolution (Fig. 3). As determined by Land (*8*), a roughly parabolic radial RI gradient across the cone is observed, with a maximum (*n* = 1.52) decreasing to *n* = 1.47 at the edge of the cone (Fig. 3A, B and C), while the RI in the intercone region drops to *n* = 1.44, due to the presence of medium-filled pore-canals (see note in SI). Furthermore, we observe a longitudinal gradient with increasing RI towards the cone tip (Fig. 3C).

**Figure 3:**
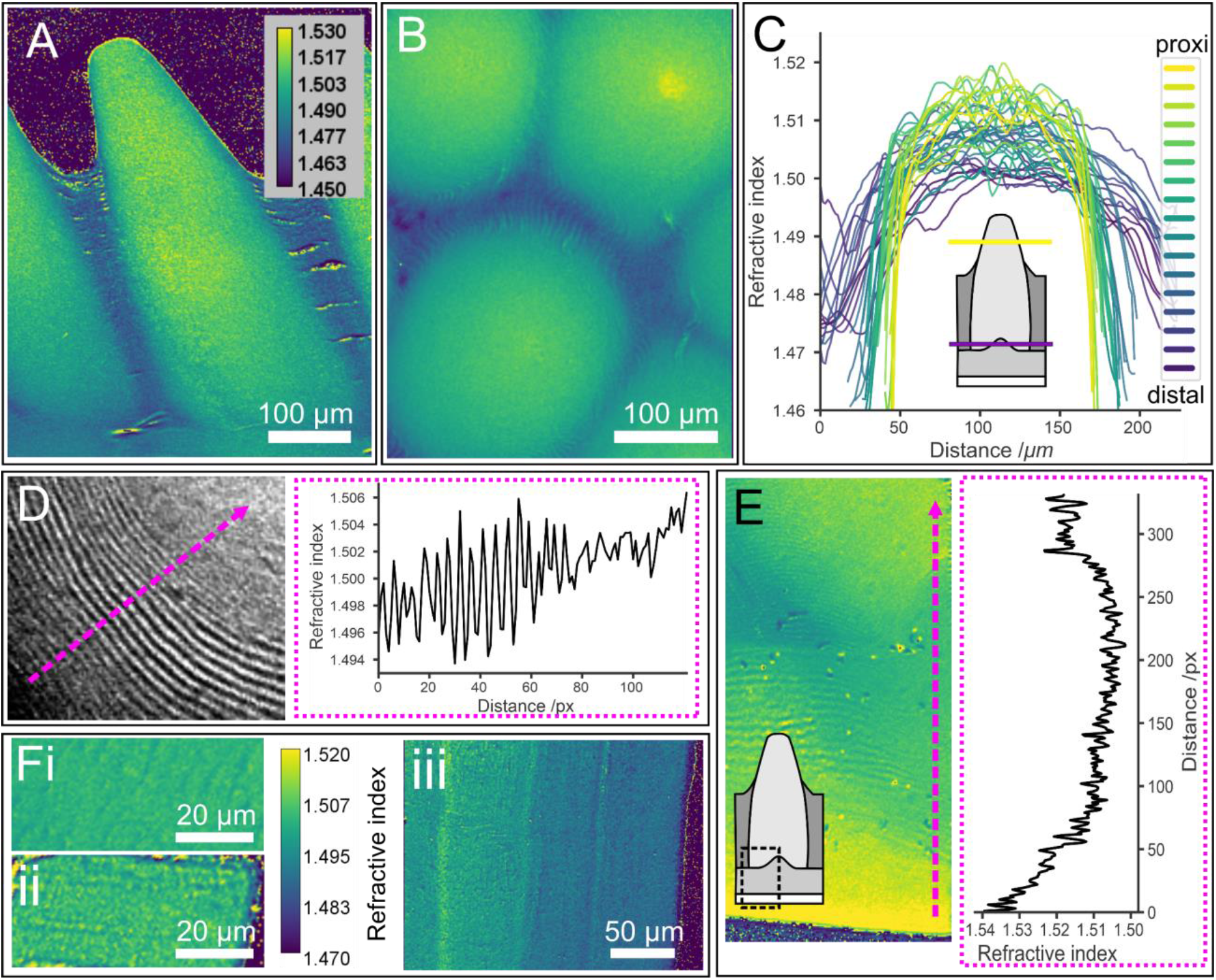
RI mapping of the cornea. RI maps are calculated from phase maps of a **(A)** longitudinal and **(B)** cross section (Same LUT as in A). **(C)** RI profiles of cross sections extracted from the base of the cone to approximately 50 μm from the cone tip. Note that the exact height in the cone cannot be determined. The profiles shown were taken from multiple cones in close vicinity to each other within single cornea **(D)** Structural RI variation observed in high magnification RI map of a cross section. The dotted magenta arrow shows the line from which the profile (mageta frame) is taken **(E)** High magnification RI map of a cornea longitudinal section around the outer-cornea protrusion (The exact area is depicted in the scheme). Note the correlation with the Br content in different cornea regions shown in Fig. 2B **(F)** RI maps of longitudinal (i) and cross (ii) sections of bleached tendons (almost pure chitin) showing similar RI value (iii) a longitudinal section of native tendon (with native proteins) demonstrating the effect of proteins on RI.

Interestingly, the maximum RI determined from QPI analysis was systematically slightly increased in cross sections relative to longitudinal sections (Fig. S3A), attesting to a structural effect caused by the helicoidal arrangement of birefringent fibers and their changing inclination across the cone. Indeed, oscillating RI values (with *Δn* = 0.01) are observed in RI maps obtained at high magnification that coincide with the lamellar texture of the cones (Fig. 3D).

Chitin is birefringent due to its orthorhombic crystal structure. Quantitative measurements and modeling of the chitin birefringence have yielded only *Δn* = 0.0024 (at 589 nm) (*27*). Indeed, the measured RI of oriented chitin fibers along and across their fiber axis, using deproteinized *L. polyphemus* tendons, show similar RI (*n* = 1.503 ± 0.005, at 650 nm), within the instrumental resolution, in both directions (Fig. 3F). In comparison, intact tendons in which the chitin crystals are decorated by ordered proteins, reveal an RI inhomogeneity that exceed the observed birefringence in the cornea (Fig. 3Fiii). These measurements demonstrate that the protein content and composition can drastically alter the material’s RI. The birefringence of ordered fibrous proteins with predominant beta-sheet structures such as silks lies in the range of Δ*n* = 0.02 to 0.04 (*28*, *29*), more than one order of magnitude higher than that of chitin. We thus attribute the structural effect of fiber orientation on the RI to ordered proteins rather than to chitin itself.

The gradient in RI cannot be explained by the structural effect alone. The higher volume fraction of proteins in the center of the cones as determined by XRD and FTIR lead to an increase in RI, while the increased water content at their periphery lowers the RI. Additionally, a sharp increase in RI (Δ*n* ~ 0.01) is observed in RI maps taken at the outer-cornea protrusion that correlate with a steep increase in BrKα XRF signal (Fig. 3E), and the maximum determined RI (*n* = 1.54) correlates with the highest level of Br at the epicuticle. Halogens in general and Br specifically have been used as dopant to increase the RI of organic polymers (*30*). This suggests that the incorporation of halogens into proteins, which is not uncommon in cuticular materials (*31*) and has been reported previously for *L. polyphemus* cuticle (*32*), may serve here to locally increase the material’s electron density and thus its RI.

We used modeling and simulations to systematically determine the role of the RI profiles and cone shape in guiding incident light into the receptor cells. Finite-difference time-domain (FDTD) simulations using a cone shape extracted from μCT data and a uniform cone RI (n = 1.52), revealed that TIR at the cone tip is sufficient to guide on-axis light to an aberrated focal point at the cone tip (Fig. 4Ai). Introducing the observed radial RI gradient drastically improves the focusing of on-axis light and places the focal point 30 μm away from the cone tip, to a point that coincides with the distal end of the receptor (*6*) (Fig. 4Aii). Implementing other anatomical details with RI variation (outer-cornea, epicuticle, and the outer-cornea protrusion) lead to negligible variation in the optical behavior (Fig. 4Aiii).

**Figure 4:**
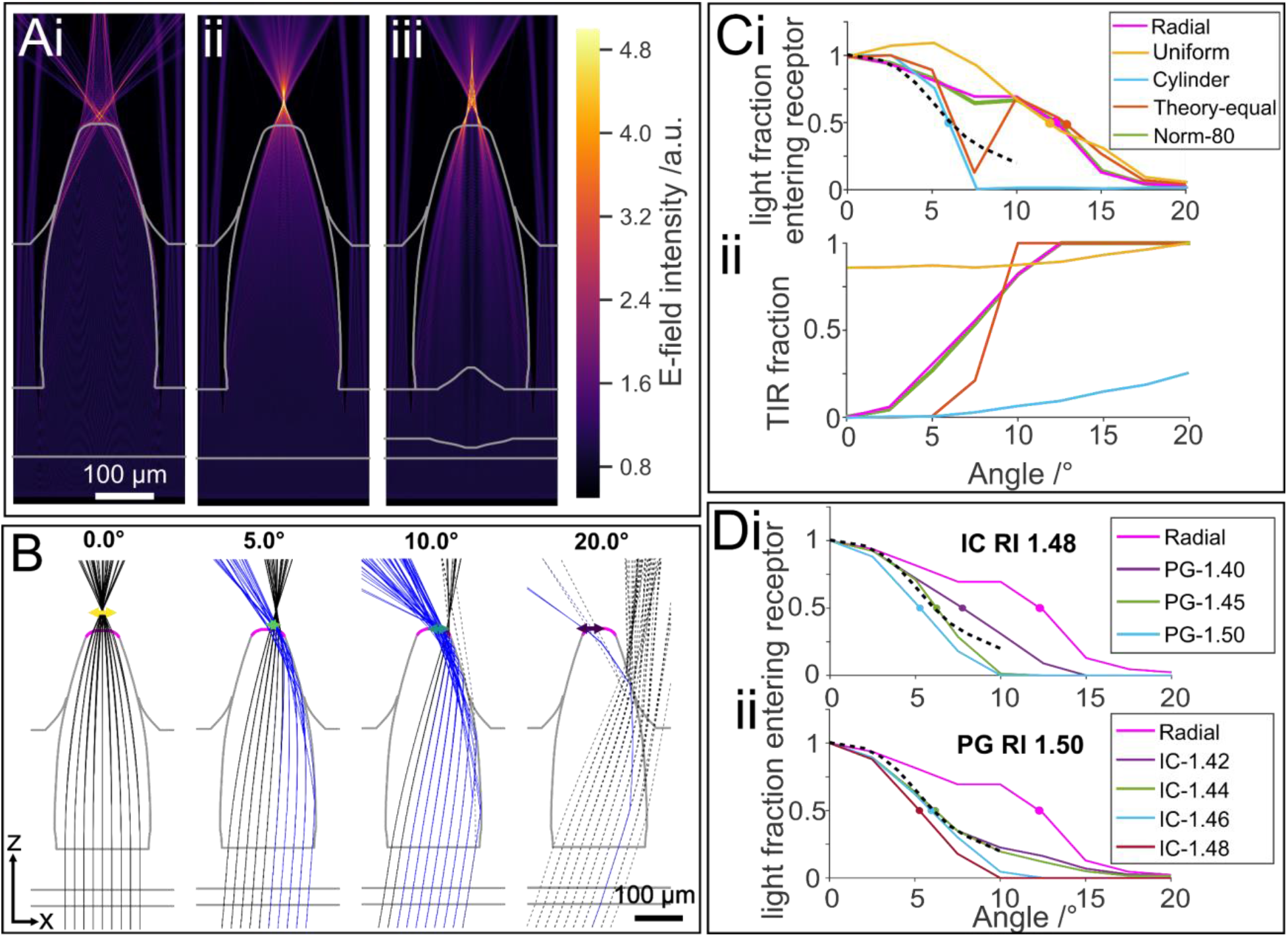
FDTD and 3D ray-tracing simulations. **(A)** FTDT simulations with 650 nm light: (i) cone model with uniform RI of 1.52, inner medium RI of 1.34 and intercone RI of 1.4, (ii) cone with the observed radial RI profile, and (iii) cone model including contributions from all subregions using the RI profile in ii. (**B)** Ray tracing results projected onto xz-plane for different incident angles with the observed RI profile and intercone value of 1.44 (pink plot in C and D). Rays that undergo TIR are plotted in blue, colored horizontal arrows represent the circle of least confusion (COLC). Rays that exit the cone before its tip (pink line) are dashed (**C)** (i) Acceptance functions based on dark adapted receptor dimensions (inner medium RI: *n* = 1.34, intercone RI: *n* = 1.40). The dashed line represents the physiological results measured by Barlow et al. 1980 (*31*). Full circles indicate the 50 % value on each curve, which represent the half-width at half-maximum acceptance angle (ii) Fraction of rays that undergo TIR before exiting the cone tip. **(D)** Acceptance functions based on dark adapted receptor dimensions with a fixed RI value for either the (i) intercone (IC) or (ii) inner medium (PG).

Ray-tracing simulations corresponded with FDTD results at normal incidence (Fig. 4B), varying the incident angle in ray-tracing simulations showed considerable variation in off-axis light focusing performance between different models. In particular, a simulated lens cylinder has a much narrower acceptance angle (Fig. 4Ci, half-width at half-maximum - 5.9°) compared to models using a cone shape, which have acceptance angles around 13°. Although the former is a close match to previous physiological measurements (6.5° (*33*)), the cylinders acceptance function has a non-physiological abrupt cut-off at higher angles of incidence. Imposing a cone shape on-top of the RI profile of the idealized lens cylinder broadened its acceptance function considerably although, surprisingly, a distinctive notch remained at 7.5° (Fig. 4Ci). The notch in the acceptance angle function disappears when the radial RI gradient profile is normalized to the changing cone diameter (Fig. 4Ci, green line), as seen in the biological system.

Examining the fraction of rays exiting the cone after undergoing TIR shows that the proportion of TIR rays tends to increase at higher angles of incidence for all models (Fig. 4Cii). To test the influence of TIR on the acceptance angle, we explored the effect of increasing the RI values of the inner medium covering the exposed cone (which would be representative of pigment granules in the distal pigment cells) and the intercone. Increasing the RI around the exposed cone tip substantially narrowed the acceptance function of the graded RI models (Fig. S5A) and could provide a closer match to the experimentally measured acceptance function for the biological relevant gradient (radial model) (Fig. 4Di). Although changing the intercone RI had little effect on either model (Fig. S5B), when the inner medium was set to 1.34, changing the intercone RI at high inner medium RI (*n* = 1.50), allowed us to find conditions (intercone RI of 1.42 to 1.44, Fig. 4Dii) that closely match the physiologically measured acceptance function.

### Conclusions

Here we have addressed the structure-property-function relationships in the *L. polyphemus* cornea by correlating spatially resolved quantitative RI measurements with structural and compositional analysis on multiple length scales. We show that the observed RI gradients have both a structural as well as a compositional origin. Most importantly, our results point toward the contribution of ordered proteins which decorate the fibrillar chitin scaffold leading to enhanced birefringence (Δ*n* = 0.01). These proteins are co-ordered with the chitin fibers and thus adopt the helicoidal arrangement. The nested arrangement of the helicoidal layers with increasing lamella inclination from the center to the edge of the cone, in turn leads to the observed radial structure-based RI gradient. The RI gradient is enhanced by compositional variations that include Br doping, a gradient in chitin to protein volume fraction and in hydration level, the latter governed by sclerotization. Sclerotization and associated cross-linking is well known for its role in stiffening the cuticle, here however, this toolkit is opted to tune the material’s optical properties. The radial RI gradient is needed for on-axis light focusing into the rhabdom. Water also plays a key role in determining the RI in the intercone region, where bulk water is present in the multiple pore-canals that span the region leading to the RI contrast that enables TIR of incoming light at higher incident angles while ensuring structural integrity. Interestingly, although the cone shape seems optimal for TIR as discussed by Levi-Setti et al. (*9*), thereby increasing the acceptance angle, this is however suppressed by the presence of screening pigment. The optical performance of the cornea is therefore guided by different mechanisms at different hierarchical length-scales. The *L. polyphemus* cornea is indeed a fascinating example of the flexibility and multifunctionality of arthropod cuticle, made with a simple “blue-print” and modular building blocks.

## Acknowledgements

This work was funded by the Deutsche Forschungsgemeinschaft under the grant no. 286895536. The authors thank the HZB and ESRF for beamtime allocation at the beamlines mySpot and ID13, respectively. Special thanks are given to Chenghao Li (MPI-CI, Department of Biomaterials), Manfred Burghammer (ESRF, ID13) and Peng Li (ESRF, ID13) for beamtime support. We are grateful to Daniel Werner (MPI-CI, Department of Biomaterials) for support with the μCT setup, Antje Völkel and Agata Baryzewska (MPI-CI, Department of Colloid Chemistry) are acknowledged for help with finding and using the Abbe refractometer. Carolin Fischer (TUD, B CUBE), Susann Weiche, Susanne Kretschmar and Thomas Kuhn (TUD, Histology facility) are being thanked for sectioning and Mallory staining. We thank Alice Ludewig (TUD, B CUBE) for support with FTIR measurements. We thank Moritz Kreysing (MPI-CBG) for useful discussion and Aurimas Narkevicius for critical review of the manuscript.

## Author contributions

Conceptualization: YP, GS, OS, PF. Project administration: YP, GS, PF, WW. Experiments performed: XRD/XRF: OS, ES, MS, JL, QPI: OS, MAKAR, TS. Raman: CS, OS. Data analysis: OS, GT, LB, YP. Modeling and simulations: FDTD: BW. Ray tracing: GT. YP, OS, GT, BW wrote the manuscript with contribution from all other authors.

## Data and materials availability

All raw data and code used for analysis are available upon reasonable request

## Supplementary Materials

Materials and Methods

Figs. S1 to S5

References (1-15)

