## Supplementary Materials for "Gradients of orientation, composition and hydration of proteins for efficient light collection by the cornea of the horseshoe crab"

Materials and Methods

Figs. S1 to S5

References (1-15)

**Materials & Methods:**

Sample preparation

*Limulus polyphemus* specimen were obtained from the Marine Biological Laboratory (Woods Hole, MA, USA), sacrificed and stored at -20 °C until use. Animals were then partially thawed at 8 °C and the lateral eyes were excised. Cells and pigments were carefully removed using fine tweezers. The remaining soft material was washed off using water spray from a syringe and a soft brush. The cleaned cornea specimens were sterilized in 70 % EtOH for 24 h and transferred to dd$H_2O$.

Large pieces of the cornea were placed in a silicone mold and covered in O.C.T medium (VWR Chemicals, Radnor, PA, USA). The samples were frozen within the cryo-microtome (560 CryoStar Cryostat, Thermo Fisher Scientific, Waltham, MA, USA) and 5 µm to 50 µm thick section were cut using Surgipath DB80 LX blades (Leica, Wetzlar, Germany) (sample −15 °C, blade −11 °C). The sections were transferred to a glass slide using a brush, thawed and rinsed 3x with dd$H_2O$ to remove excess O.C.T. Section samples were used for Raman micro-spectroscopy, confocal laser scanning microscopy, QPI, FTIR mapping and XRD/XRF.

Thin sections for Mallory staining were prepared at the CMCB-EM and histology facility. Cornea pieces were embedded in epon, sliced 1 µm to 2 µm thick using an ultramicrotome and mounted on glass slides.

Confocal laser scanning microscopy (CLSM)

To investigate the chitin fiber architecture, we used 5 µm sections of cornea dyed with 0.4 mg/mL DY96 (Sigma-Aldrich, St. Louis, MO, USA) in water. DY96, which binds to chitin and other polysaccharides (*1*), only emits a detectable fluorescence signal when the chitin fibers are parallel with the imaging plane (*2*). Imaging data was acquired with a SP-8 laser scanning microscope (Leica, Wetzlar, Germany) equipped with a 63x water objective (NA=1.2). A multi photon laser (Spectra-Physics, Stahnsdorf, Germany) at 780 nm was used for excitation and the signal was recorded on a HyD detector between 540 nm and 580 nm.

Micro Computer Tomography (µCT)

µCT was performed on hydrated cornea portions stained with iodine vapor for 12 h. The stained samples were placed in a plastic vial and kept in contact with water to allow full hydration of the sample during the measurement. Transmission scans were acquired using an EasyTom160/150 equipped with a micro-focus X-ray source using a tungsten filament and a digital flat panel detector (RX Solutions, Chavanod, France). The acceleration voltage was set to 70 kV, with a current of 100 µA recording 1120 images at full rotation with a total measurement time of 3.7 h at a sample-to-detector distance of 773.193 mm yielding a voxel size of $(2.18\ \mu m)^3$. Reconstruction of the final volume was performed using the X-Act software (RX Solutions, Chavanod, France).

Fourier transform infrared spectropscopy (FTIR)

FTIR data was acquired using a Lumos II micro-spectrometer equipped with a nitrogen-cooled mercury cadmium telluride detector in transmission (Bruker, Billerica, MA, USA). To reduce noise from water vapour and $CO_2$, a chamber with continuous $N_2$ was placed around the microscope stage. Thin cornea sections of 5 µm were placed on $CaF_2$ windows (Korth Kristalle GmbH, Altenholz, Germany) for measurements. For background correction, spectra of the sample carrier were acquired by averaging 32 interferograms with a step size of 4 $cm^{-1}$ each time before measuring the sample. Subsequently the transmittance spectrum of the sample was recorded in the same manner. For measurement control and data analysis Opus spectroscopy v8.5.29 software was used. The transmission spectra were converted into absorbance spectra and the ratio of the peaks between 1700 $cm^{-1}$ to 1600 $cm^{-1}$ and 1180 $cm^{-1}$ to 1000 $cm^{-1}$ (Fig. S3C).

Raman microspectroscopy

Raman microspectroscopy was performed on 10 µm hydrated cryo sections mounted between a glass slide and a cover slip and sealed with nail polish. We used an alpha300 confocal Raman microscope (WITec, Karlsruhe, Germany) equipped with a P-500 piezo-scanner (Physik Instrumente, Karlsruhe, Germany). The laser ($\lambda$ = 532 nm) was focused by a 40x objective lens (NA = 1.2, Nikon, Tokyo, Japan) with a laser power of 15 mW on the sample. Spectra were acquired by a thermoelectrically cooled CCD detector (DU401A-BV, Andor, Belfast, North Ireland) behind a 600 g $mm^{-1}$ grating spectrograph (UHTS 300, WITec, Ulm, Germany) with a spectral resolution of 3 $cm^{-1}$. The WITec Control software (Version 5.2, WITec, Ulm, Germany) was used for measurement setup and control. The hydration map was generated using the WITec Project Five v5.2 software by mapping the integrated intensity of the OH bands between 3.111 $cm^{-1}$ and 3.689 $cm^{-1}$ (*3*) (Fig. S3D).

Mallory trichrome staining

For Mallory staining microtome sections of 1 µm and 2 µm thickness were created from epon-embedded cornea pieces and transferred (using 10 % acetone) to glass slides which were dried over night at 40 °C. Sections were incubated 30 min in NaOH/100 % EtOH to etch the epon followed by 3x washing in 96 % EtOH and 10 min under running $ddH_2O$. This was followed by 15 min at 60 °C in etchant (1:1 potassium dichromat in $ddH_2O$ and 10 % HCl in 96 % EtOH) and 10 min under running tap water. After 1 min in $ddH_2O$ the Mallory trichrome staining (1g phosphotungstic acid, 2 g Orange G, 1 g aniline blue, 3g fuchsin acid in 200 mL $ddH_2O$) was applied for 15 min at 60 °C. After brief washing in $ddH_2O$ the glass slides containing the sections were subsequently briefly bathed in 70 % EtOH and 3x in 96 % EtOH. Finally, the sections were immersed in 100 % EtOH for a maximum of 1 min, followed by two 2 min immersions in xylol. The samples were then dried and sealed with Cytoseal.

Refractive index mapping

RI mapping was performed using a Q-Phase microscope (TELIGHT, Brno, Czech Republic), employing quantitative phase imaging (QPI) based on holographic microscope with low-coherent illumination (*4*). During a measurement, the phase shift caused by the sample is retrieved, and used to calculate the sample RI using the relation:

$$n_s = \frac{\varphi\lambda}{2\pi h_s} + n_m, \qquad \text{(S1)}$$

where $n_s$ is the sample effective RI, $h_s$ the local sample thickness, $\varphi$ the measured phase shift, $\lambda$ the light wavelength (here 650 nm) and $n_m$ refractive index of the surrounding medium. In order to overcome uncertainties related to sample thickness, the measurements were carried out using two media with a different RI. The RI of the sample can then be calculated using the following equation derived from equation 1 (*5*):

$$n_s = \frac{\varphi_2 n_{m1} - \varphi_1 n_{m2}}{\varphi_2 - \varphi_1}, \qquad (S2)$$

where indices 1 and 2 refer to the two different media used.

Sections of 5 µm thickness were glued to a glass slide using a small drop of UHU max repair (UHU, Baden-Baden, Germany) to prevent movement during medium exchange and covered by a sticky-Slide (ibidi GmbH, Gräfelfing, Germany) with a 200 µm high channel. To change the medium RI we used Optiprep (60 % Iodixanol in water, n=1.429 at 591 nm (*6*), Stemcell Technologies, Köln, Germany) at different dilutions. The RI (at 650 nm) of the iodixanol solutions was determined using an Abbe refractometer with a white light source and a 650 nm filter. During QPI measurements, medium exchange was performed using a syringe connected by Luer connectors and silicone tubing to the perfusion slide. The sample chamber and connections were sealed using parafilm. After obtaining the phase map at different ROI, the channel was flushed with 3 mL of the new solution to ensure complete exchange of medium before continuation of measurements. Data analysis was performed using in-house python scripts and CV2 libraries for alignment of the images and calculation of RI maps.

In order to define the cornea RI gradients, we selected a data set extracted profile plots (Fiji (*7*)) from cornea cross sections obtained at various heights along the length of cones from subsequent sections of a single animal, imaged using 4x objective. The profiles were ordered from distal to proximal depending on section number and cone diameter. The comparison of maximum RI of longitudinal and cross cornea sections (Fig. S3A) was performed on sections from cones in close proximity from a single animal mounted on the same slide. Pairs of maximum RI values were created based on the height along the cone of the respective cross section and the height of the profile taken on the longitudinal section.

RI of intercone: The phase-decoupling method provides a quantitative measure of the solid material but effectively disregards free medium. The measured values for the intercone region containing water-filled pore-canals were corrected based on the pore canal density estimated by thresholding CLSM images of cornea cross sections (Fig. S1 C). Pore canal volume fraction was estimate by measuring the ratio of masked pixels to total area. This resulted in pore canal density around 15 %. Assuming RI of 1.34 for the pore canal filling medium, the final RI of the intercone regions obtained is 1.44.

X-ray diffraction (XRD) and X-ray fluorescence (XRF)

Cone sections of 50 µm thickness, prepared using a cryo-microtome (see above), mounted on SiN windows, were measured at mySPOT beamline at BESSY II, HZB (Helmholtz-Zentrum Berlin für Materialien und Energie, Berlin, Germany) and at ID13 beamline at ESRF (European Synchrotron Radiation Facility, Grenoble, France). In both cases the beam energy was set to 15 keV (0.827 Å), a diffraction detector in transmission geometry with a sample-to-detector distance of around 300 mm and fluorescence detector at roughly 90° relative to the beam was used.

At the mySpot beamline the beam energy was set with a B4C/Mo Multilayer monochromator and an Eiger 9M (pixel size (75x75) µm², Dectris, Baden, Switzerland) XRD detector was used with a (30 µm)² beam. Quartz was used for detector calibration and a single element Si(Li) XRF detector (RAYSPEC (SGX Sensortech) was used for XRF. For hydrated maps, sections in water were placed in-between SiN windows sealed with vacuum grease to prevent evaporation. Oversampling with step size of 8 µm was used in both directions at an acquisition time of 10 s.

At the ID13 beamline the beam energy was set with a Si(111) monochromator and an Eiger 4M (pixel size (75 µm)², Dectris, Baden, Switzerland) XRD detector was used with a focal size of (300 nm)². Quartz and silver behenate powders were used for detector calibration. A Vortex EM, silicon drift detector with 50 mm² sensitive area was used for recording the XRF signal. Maps were acquired with a step size of 1 µm in x and y and an acquisition time of 100 ms.

XRF maps were generated from integrating the BrKα peak intensity and then corrected by primary beam intensity. XRD data reduction was performed using dpdak (*8*) v1.4.1 to obtain XRD 1D profiles. Background and primary beam correction were performed using in-house python scripts based on the numpy library. Fiber correlation peak fitting was performed using dpdak. Protein/chitin volume fractions and water content were calculated using equation 1 (*9*). The profiles shown in Fig. 2 were averaged over ~100 diffraction patterns in the corresponding region using in-house python scripts using sympy library.

Water volume fraction: The calculation of the water volume fraction follows the assumption that bound water is retained in the air-dried section, and the total volume is a sum of chitin and proteins. In the hydrated samples the total volume consists of chitin, proteins and water. We assume that the ratio of chitin and proteins is unchanged, and calculate the hydrated protein fraction from the hydrated chitin fraction multiplied by this ratio (equation 2).

Chitin crystallite size: The in-plane fiber orientation can be retrieved from the orientation of the correlation peak on the 2D detector, readily confirming the nested organization of the helicoidal lamellae seen with CLSM (Fig. S2 A). To confirm that chitin crystallize size is uniform across the cornea, we fitted averaged XRD spectra from different regions across the cone. For this, the (110) peak was fitted with four Gaussians to account for the additional signal from other chitin and protein reflections (Fig. S2 B). The fitting parameters of the main peak were used to solve the Scherrer equation

$$\tau = \frac{K * \lambda}{\beta * \cos(\theta)}, \qquad (S3)$$

where $\tau$ is the crystallite thickness, K is a form factor (here set to 1), $\lambda$ the wavelength used (0.82656 Å), $\beta$ the FWHM and $\theta$ the scattering angle of the reflection. The average result for chitin crystallite thickness from 14 different regions within the cornea was 24.52 Å $\pm$ 2.37 Å. Assuming a distance of b = 18.9 Å (*10*) for the respective crystallographic plane, this leads to a chitin fiber diameter of 28.35 Å or 1.5 unit-cells in this direction. This slight offset is most likely due to the large number of degrees of freedom resulting from the simultaneous fitting of four peaks.

Model generation for FDTD and ray tracing analyses

The µCT volume data was segmented using Amira (v5.3) to label the extent of six cones, as well as the surrounding intercone, outer cornea, and epicornea. A Matlab script was developed to calculate the radially averaged surface profile along each cone, as well as its outer-cornea protrusion, epicornea dent, and the bordering of internally exposed intercone. The angle between the central axis of the cone and corneal normal differed by 22.2° $\pm$ 1.5° in the section of the cornea that was scanned. To remove the influence of off-axis optics from the analysis we aligned the surfaces of the outer cornea and epicornea to each cone by rotating the corneal normal to align with its central axis and then translated the tip of the cone-in-cone and epicornea cone so that they were on the axis. The internally exposed intercone surface lies in parallel to the cornea and was also rotated to align it with the central axis of each cone, however, as the intercone surface also follows the cone shape, a shear transformation then had to be applied to the intercone so that its alignment remained consistent with the cone surface after rotation. Finally, the outer-cornea protrusion and epicuticle dent were shifted such that the cone axis passed through their center.

As the length of each of the six analyzed cones varied slightly (215.4 µm $\pm$ 7.4 µm), the radial outlines of each structure were stretched to a uniform length before calculating an average radial outline, which was stretched back to the average cone length (Fig. S5C), and finally smoothed. The averaged radial profile of each structure was used to generate either 2D RI maps at (500 nm)$^2$ resolution for FDTD simulations or 3D RI maps at (2 µm)$^3$ resolution for ray-tracing simulations. Triangulated meshes were also calculated to delineate the border of each surface. Unless otherwise noted, the RI map used the following values for each structure: outer media – 1.33; epicornea – 1.53; outer cornea – 1.5; intercone – 1.40; cone – 1.52; inner media – 1.34. In addition to using a constant RI value for the cone, we also modelled the following RI profiles:

Observed: $RI(r,z) = 1.5 + (z \times 0.01 + 0.01) - (0.5 \times z + 0.8) \times 0.0392 \times r^2$, where *z* is the vertical distance from the cone tip (normalized to 1) and *r* is the radial distance from cone center line (normalized to 1, relative the radius at each step along the profile).
The linear component of the observed profile could be modelled separately by setting *r* = 0. Likewise, the radial component could be modelled separately by setting *z* = 0 or 1 for the cone tip or base, respectively (Fig. S4A).

Theoretical: i.e. the parabolic RI profile required to focus light at the tip of a lens cylinder of arbitrary length (*11*). The profile was defined as: $RI(r) = 1.52 \times \mathrm{sech}(\frac{\pi r}{l})$, where *r* is the absolute radial distance from the cone center line and *l* is the absolute length of the cone. Besides modelling a cone with this theoretical RI profile, we also modelled a full cylinder, with a radius equal to the maximum of the average radial cone profile.

Ray-tracing

3D ray-tracing was performed by implementing the simulation procedure for media with discretely defined isotropic but inhomogeneous refractive index developed by Nishidate et al. (*12*) in a Matlab script. The following two modifications were made to the procedure published by Nishidate et al.: 1) Numerical integration was performed using the 4$^{th}$ order Runge-Kutta scheme described by Sharma et al. (*13*) with a fixed step size of 0.5 µm, rather than the Runge-Kutta-Fehlberg scheme with an adaptive step size; 2) refraction at lens surfaces were calculated based on the ray's intersection to the triangulated mesh of the lens surface, rather than using a Nagata patch. Our implementation achieved comparable accuracy to that reported by Nishidate et al. (*12*) when tested on a Luneburg lens and an optical fiber with a parabolically graded RI. Simulations were conducted for rays placed at 10 µm intervals on two-dimensional grid and for angles (relative to the cone axis) from 0° to 20° spaced by 2.5° intervals. Rays that did not enter the base of the cone (i.e. rays that did not enter the cone or entered along its side) were not included in the analysis, nor were rays that reversed direction due to total internal reflection (TIR).
Ray-tracing results were described by plotting orthogonal perspectives of ray paths (paths are shown for rays at 20 µm intervals [i.e. every 2$^{nd}$ ray simulated] to preserve readability) and spot diagrams of the position of all rays on a plane passing through the cone tip. Anatomical studies indicate that the cone is sheathed in pigment until approximately 15 µm away from the cone tip, while the top of the receptor is positioned 5 µm below the cone tip and has a radius of 25 µm in dark-adapted animals at night (*14*) (see the top diagram in Fig. S4B). As such, we calculated the height and radius of the best focus in both perspectives as well as for the circle of least confusion

(COLC), from rays that exited the cone less than 15 µm from its tip. The approximate receptor acceptance function was calculated as the number of rays that exited the cone tip and then entered the receptor within its dark-adapted radius for each angle of incidence (which was normalized to a fraction based on the number of rays entering the receptor at 0°). This approximation represents an upper bound on an acceptance function, as the proportion of light absorbed from each ray by the receptor will vary depending on the rays entry position and angle. A plot of relative sensitivity as a function of illumination angle (for night adapted receptors) measured from optic nerve fibers by Barlow et al. (*15*) (Fig 4C) was digitized using WebPlotDigitizer and averaged around 0°; this provides an independent reference for our acceptance function calculations. We also calculated an acceptance function based on the day-adapted receptor dimensions and pigment aperture (see the bottom diagram in Fig. S4B), and found that light only reached the receptor at normal incidence. This is a limitation caused by the resolution used in the ray-tracing model, as the half-width measured for the physiological light-adapted acceptance function is 3$^{o}$ (*15*).

Finite-difference time-domain (FDTD) simulations

Light scattering by the internal structure of the cones was simulated using a two-dimensional FDTD method in Lumerical FDTD 2012 R1.4 (Ansys Canada Ltd., Vancouver, Canada; https://www.lumerical.com/products/fdtd/), a commercial-grade FDTD simulator. We performed FDTD simulations based on the different models obtained from µCT scans as explained above. The incident light beam was assumed to be a plane-wave with a broad-band spectrum centered around 650 nm, the wavelength at which the RI maps were obtained. The electric field maps were calculated from light flux through the structure.

**Supplemental Figures**

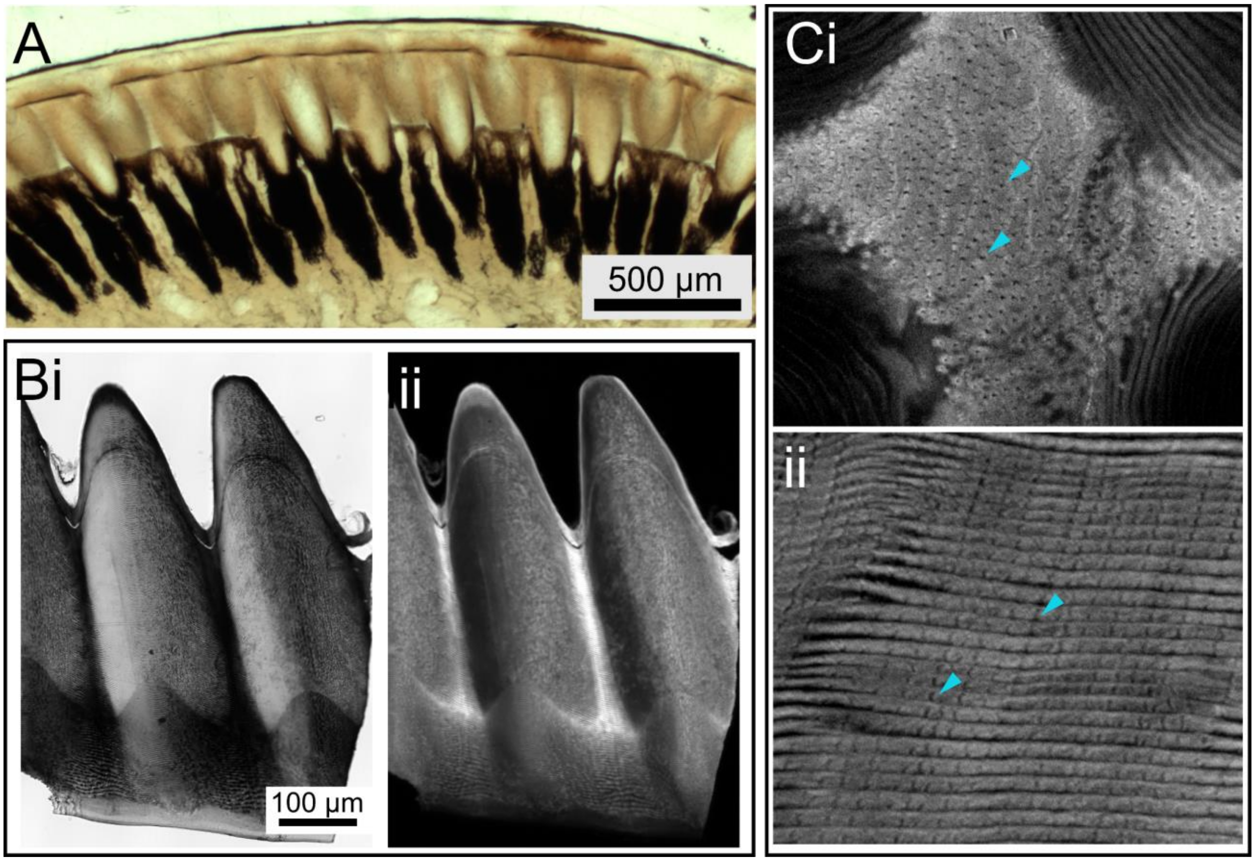


**Fig. S1. Cornea sections: lamellae and pore canals (A)** Longitudinal section through untreated cornea embedded in PMMA shows the corneal cone arrangement and the pigment enveloping the cones. The amber color results from the embedding procedure. The cornea is typically light amber and transparent. **(B)** Low magnification of cornea longitudinal section imaged using CLSM (i) bright-field image and (ii) DY96 channel. Note that the epicornea is unstained indicating the absence of chitin **(C)** Magnified intercone regions showing multiple pore canals (dark structures, two are indicated by arrows) in (i) cross section and (ii) longitudinal sections. The lamellated organization of the material in the cones (Ci) and the intercone region (Cii) is also clearly visible.

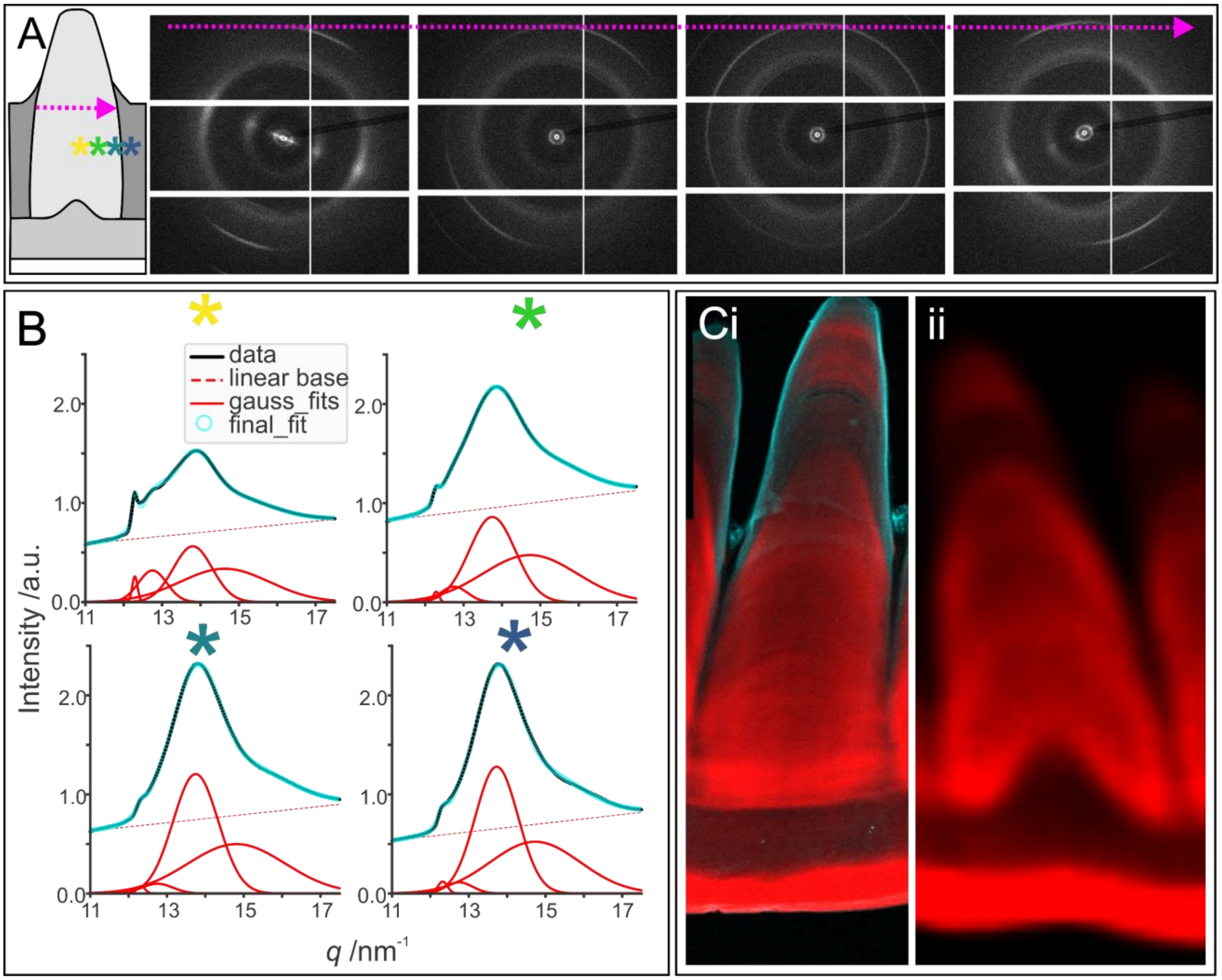


**Fig. S2. XRD/XRF analysis of cone longitudinal sections (A)** 2D diffraction patterns across a longitudinal cornea section showing the typical chitin pattern and the fibrous nature of the cuticular cornea. The rotation of the anisotropic fiber signal confirms the nested helicoidal structure observed by CLSM: the pink arrow indicates the direction across the cone from which the diffraction patterns were taken. Colored asterisks indicate the positions of plots in B **(B)** Fitting on the (110) reflection to obtain parameters for calculating the chitin crystallite thickness in the cornea by the Scherrer equation. **(C)** XRF maps of Br (red) and Zn (cyan) (i) shows the Zn layer surrounding the exposed part of the cone. The section does not contain the outer-cornea protrusion due to oblique sectioning. (ii) Low resolution map of a longitudinal section through the center of the cone, showing the distribution of Br around the outer-cornea protrusion.

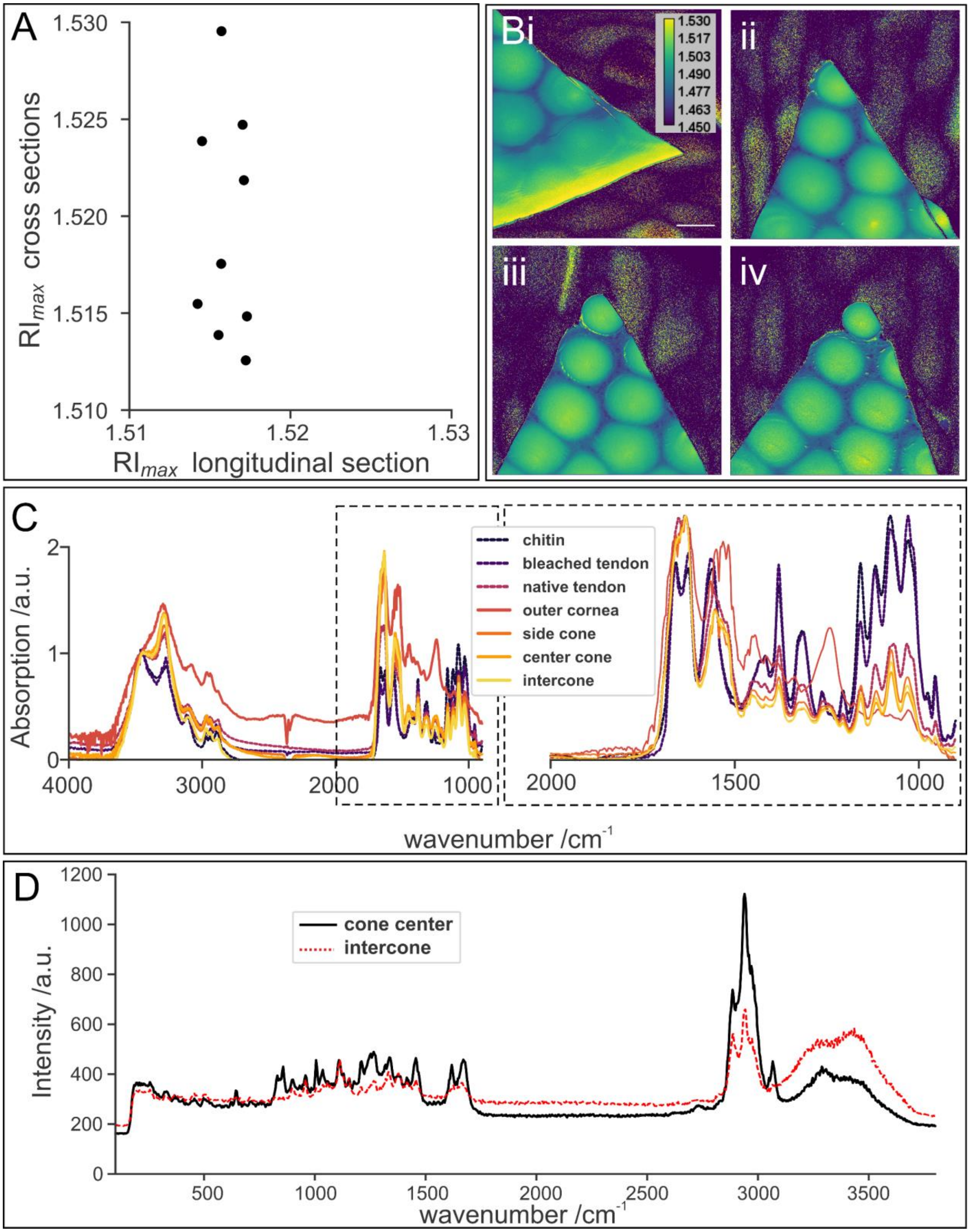


**Fig. S3. RI mapping and compositional variation (A)** Pairs of maximum RI values of cross and longitudinal sections of the same animal, sectioned from corneal cones in close proximity and measured at the same experimental session. Pairs were chosen to be at approximately the same height along the cone. **(B)** RI maps of a cornea cross section in a curved cornea region, allowing access to the outer cornea (top left) **(C)** FTIR spectra of chitin reference as well as native and deproteinized *L. polyphemus* tendons compared with FTIR spectra of different regions of the cornea. The full range is normalized to the peak at 3450 $cm^{-1}$, the spectra of the magnified plot are normalized to their respective maximum values, emphasizing the individual contribution of chitin and protein. The map in Fig. 2G shows the ratio of the integrated signal of chitin and protein over the integrated chitin intensity (1700 $cm^{-1}$ to 1600 $cm^{-1}$ over 1180 $cm^{-1}$ to 1000 $cm^{-1}$). **(D)** Raman spectra at the cone center and at the intercone. The hydration map shown in Fig. 2E was created by integrating the intensity of the OH bands between 3111 $cm^{-1}$ and 3689 $cm^{-1}$.

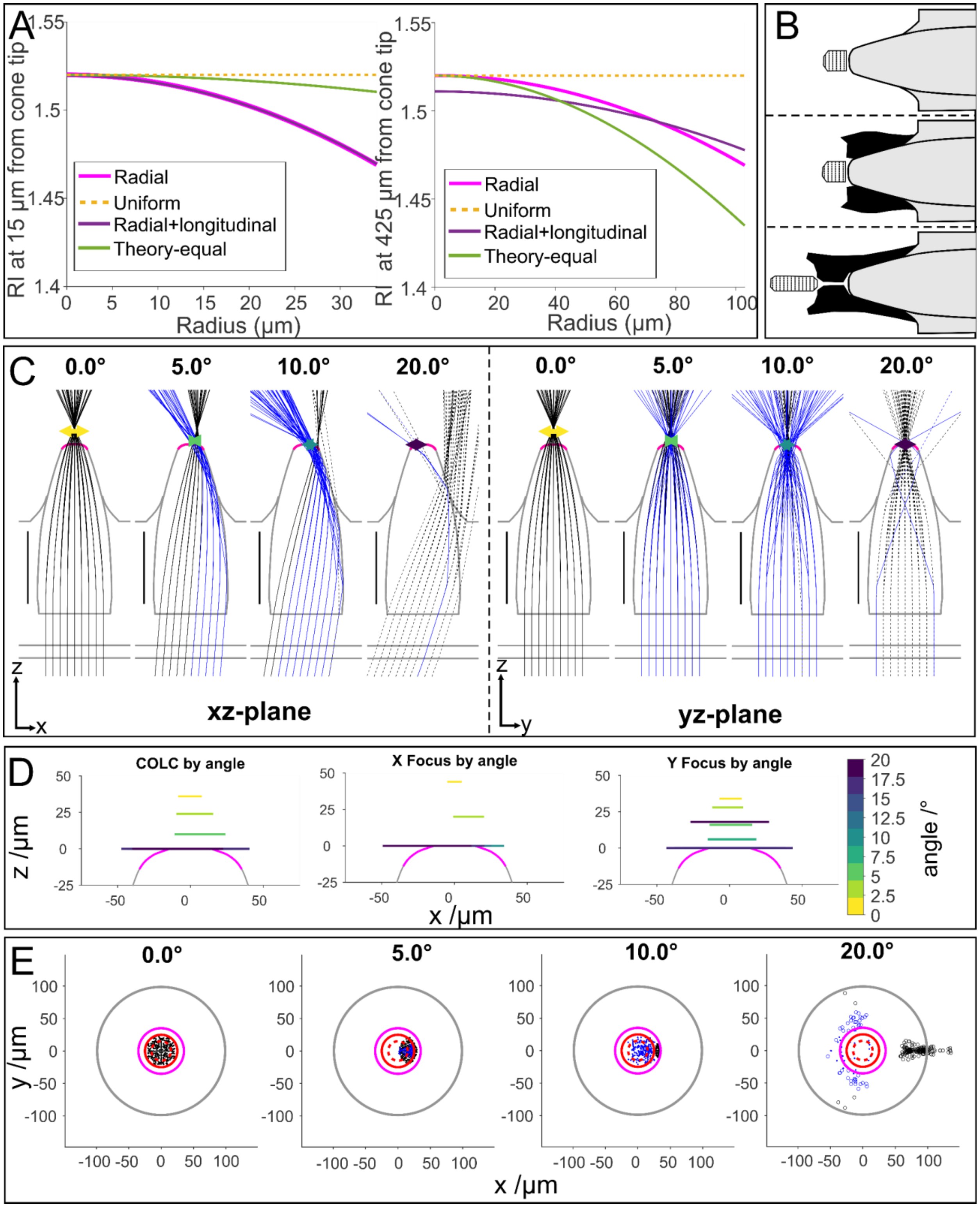


**Fig. S4. 3D Ray-tracing (A)** RI profiles used for ray tracing. The left plot is calculated at the tip of the exposed cone, the right plot is at the widest (base) part of the cone. **(B)** Receptor position and pigment (depicted in black) defining the aperture in light or dark adaptation according to

Chamberlain and Barlow 1987 (*14*) top to bottom: dark adapted model with low inner medium RI (*n*=1.34), dark adapted with pigment (n=1.50 for inner medium), light adapted with narrow elongated aperture **(C)** Ray focusing in the xz- and yz-plane on the example of 'radial' gradient model for different incident angles (the same model as shown in Fig. 4B). The pink outline on the cone tip shows the exposed cone tip (also shown on the outlines in D and E). The circle of least confusion (COLC) is depicted as a colored horizontal arrowed bar, rays that undergo TIR are plotted in blue (also shown in E), and rays that exit the cone before its tip are dashed (open circles in E). **(D)** Same model as in C showing lines denoting the position and diameter of the COLC, X focus and Y focus for all angles simulated (note that the focal heights usually descend to the cone tip at angles greater than 7.5°, in which case the lines overlap). The color bars denote the incidence angle in degrees. **(E)** Spot diagrams (at the cone tip) for the same cone model, showing the rays (dots) exiting the cone tip (pink circle) and entering the receptor (night – solid red circle, day – red dashed circle) for selected angles. The cone base is represented by the grey circle.

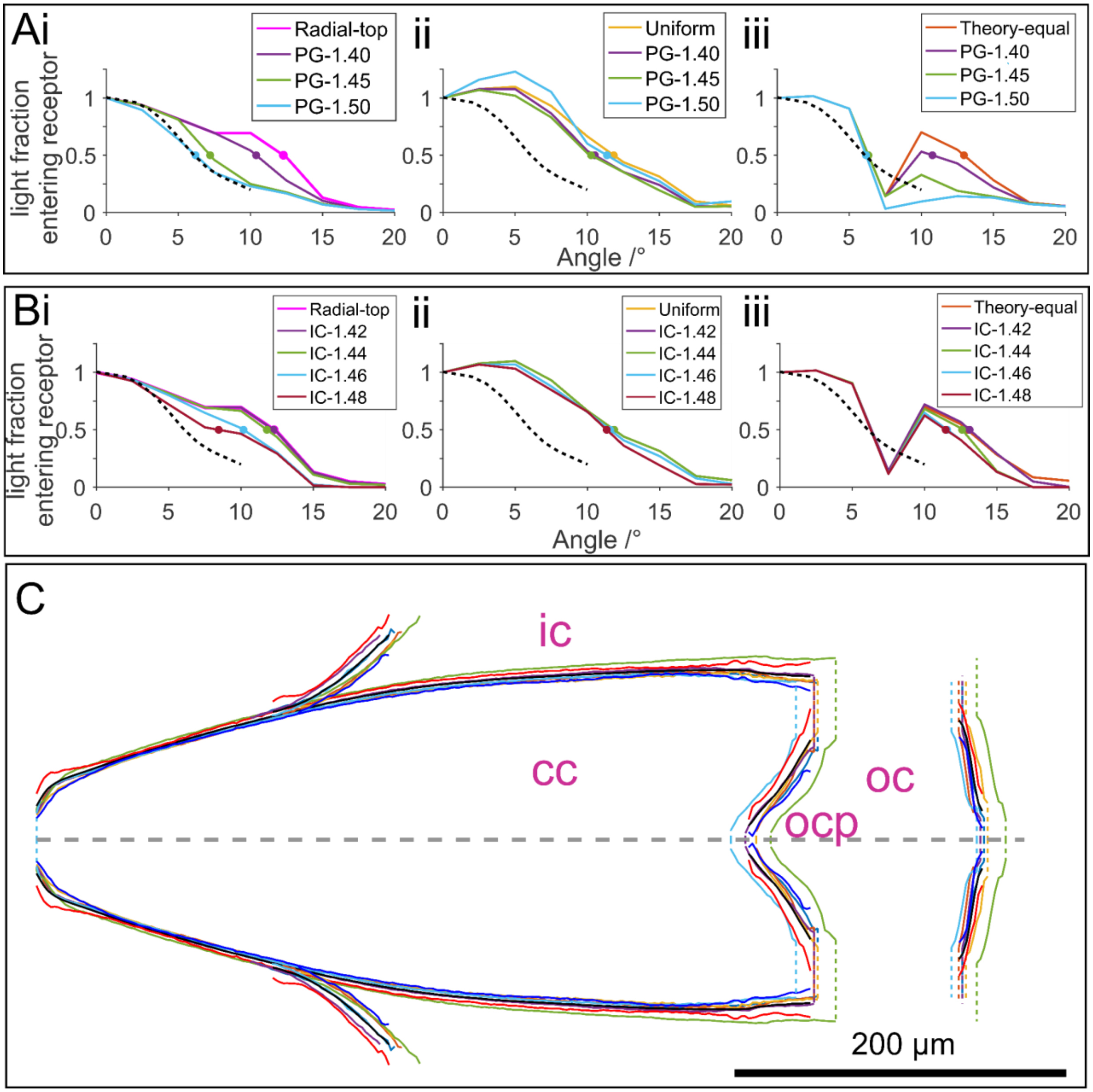


**Fig. S5. (A)** Effect of varying RI of screening pigment around exposed part of cone on acceptance function for three different models (intercone RI: *n*=1.40). **(B)** Effect of varying the RI of the intercone on acceptance function for three different models (inner medium RI: *n*=1.34). **(C)** Diagram showing variation in the cone outline measured from six segmented groups of the corneal cone (cc), outer-cornea protrusion (ocp), intercone (ic) and outer cornea (oc). Black lines indicate the averaged profile, while the blue and red lines indicate the mean minus or plus 4 SD, respectively. The dashed grey line represents the central axis (and average length) of the cone around which all profiles are symmetric.

**References (1-15)**

12. Y. Nishidate, T. Nagata, S. Y. Morita, Y. Yamagata, Ray-tracing method for isotropic

inhomogeneous refractive-index media from arbitrary discrete input. *Appl. Opt.* **50**, 5192–5199 (2011).

13. A. Sharma, D. V. Kumar, A. K. Ghatak, Tracing rays through graded-index media: a new method. *Appl. Opt.* **21**, 984-987 (1982).

14. S. C. Chamberlain, R. B. Barlow, Control of structural rhythms in the lateral eye of *Limulus*: interactions of natural lighting and circadian efferent activity. *J. Neurosci.* **7**, 2135–2144 (1987).

15. R. B. Barlow, S. C. Chamberlain, J. Z. Levinson, *Limulus* brain modulates the structure and function of the lateral eyes. *Science.* **210**, 1037–1039 (1980).
